# Motivational, proteostatic and transcriptional deficits precede synapse loss, gliosis and neurodegeneration in the B6.*Htt*^*Q111/+*^ model of Huntington’s disease

**DOI:** 10.1101/081109

**Authors:** Robert M. Bragg, Sydney R. Coffey, Rory M. Weston, Seth A. Ament, Jeffrey P. Cantle, Shawn Minnig, Cory C. Funk, Dominic D. Shuttleworth, Emily L. Woods, Bonnie R. Sullivan, Lindsey Jones, Anne Glickenhaus, John S. Anderson, Michael D. Anderson, Stephen B. Dunnett, Vanessa C. Wheeler, Marcy E. MacDonald, Simon P. Brooks, Nathan D. Price, Jeffrey B. Carroll

## Abstract

We investigated the appearance and progression of disease-relevant signs in the B6.*Htt*^*Q111/+*^ mouse, a genetically precise model of the mutation that causes Huntington’s disease (HD). We find that B6.*Htt*^*Q111/+*^ mice are healthy, show no overt signs of central or peripheral inflammation, and no gross motor impairment as late as 12 months of age. Behaviorally, we find that 4-9 month old B6.*Htt*^*Q111/+*^ mice have normal activity levels and show no clear signs of anxiety or depression, but do show clear signs of reduced motivation. The neuronal density, neuronal size, synaptic density and number of glia is normal in B6.*Htt*^*Q111/+*^ striatum, the most vulnerable brain region in HD, up to 12 months of age. Despite this preservation of the synaptic and cellular composition of the striatum, we observe clear progressive, striatal-specific, transcriptional dysregulation and accumulation of neuronal intranuclear inclusions (NIIs). Simulation studies suggest these molecular endpoints are sufficiently robust for future preclinical studies, and that B6.*Htt*^*Q111/+*^ mice are a useful tool for modeling disease-modifying or neuroprotective strategies for disease processes before the onset of overt phenotypes.

## Introduction

Huntington’s disease (HD) is an autosomal dominant neurodegenerative disease caused by expansion of a glutamine-coding CAG trinucleotide repeat near the 5’ end of the *HTT* gene ^1^. The identification of the disease-causing mutation did not point to an obvious set of therapeutic approaches, given that the huntingtin protein is a very large HEAT/HEAT-like repeat solenoid scaffold ^2^, which is highly evolutionarily conserved, and vital to normal development ^3^. A key effort in preclinical HD research is understanding the link between polyglutamine expansion in huntingtin and the selective cellular toxicity in corticostriatal circuits which is thought to cause disease symptoms. A range of animal models of the HD mutation have been developed to investigate this question, including knock-in and transgenic mice as well as transgenic rats, sheep, pigs and nonhuman primates ^4^. Unique amongst described models, knock-in mice with varying allele sizes provide the ability to quantitatively study the relationship between CAG size and pathogenesis in an otherwise isogenic system ^5^. Recent molecular ^6^ and behavioral ^7^ analyses of an allelic series of HD knock-in mice reveal very discrete CAG-dependent disease signatures, suggesting these mice recapitulate a cardinal feature of human HD, namely the relationship between CAG tract length and the rapidity of disease progression ^3,8^. Because knock-in models express *Htt* at endogenous levels from the endogenous locus they precisely mimic the genetics of human HD, however their use in preclinical studies has been limited because their overt neurological signs are very subtle compared to transgenic animals ^9,10^. The bulk of preclinical research has therefore been conducted in transgenic models expressing either full-length or short fragments of mutant huntingtin, which display more phenotypes ^11^.

The embracing of transgenic models by the field has largely been driven by the desire to model important signs and symptoms of HD, including progressive striatal atrophy and hyperkinetic motor disturbances. We consider that these phenomena emerge from a substrate of decades of progressive cellular and physiological dysfunction before manifesting in human patients as a set of clinical signs recognizable as “Huntington’s disease” ^12^. Targeting these late features of the disease process, including cell death and emergent motor symptoms, may enable the modification of late HD symptoms. However, this strategy doesn’t allow targeting of the earliest disease changes that precede overt clinical disease features, which if clearly identified might ultimately provide paths to prevention and/or early-stage reversal. Finally, no HD mouse model develops chorea-like hyperkinetic movements, and mouse models present with modest striatal volume loss over the lifespan of the mouse ^13,14^, while human disease results in the nearly complete atrophy of this critical structure ^15^.

We and others acknowledge that knock-in mouse models of HD cannot fully recapitulate clinical symptoms that, in human mutation carriers, require approximately four decades of pathogenic processes to emerge and are followed by a further 15 years to progress to death ^16^. However, these genetically faithful but subtle mouse models provide us the ability to study the earliest and subtlest changes caused by the CAG expansion of *Htt* - the very processes we wish to understand if we are to develop interventions to prevent the development of clinical HD onset, rather than addressing its symptoms only after significant damage to key processes has been incurred. Towards this end, we have here characterized the earliest observable changes in the murine B6.*Htt*^*Q111/+*^ (hereafter *Htt*^*Q111/+*^) model of the HD mutation ^17^. Since their creation, these mice have been utilized to test specific hypotheses in a wide range of studies ^18^, and a single unbiased phenotypic screen ^10^, but have not been well characterized as a system for conducting preclinical studies. We therefore examined the incidence, progression and statistical power of a range of phenotypes in these mice, and find that they provide a powerful tool for studying hypotheses about the early disease events that stem from the HD CAG expansion mutation. We observe robust transcriptional, proteostatic and motivational phenotypes which progress from 3 to 12 months of age in *Htt*^*Q111/+*^ mice. Simulation studies suggest interventions that rescue specific signs by 10-25% are readily detectable in appropriately powered preclinical studies, making the *Htt*^*Q111/+*^ mouse a useful tool for testing disease modifying therapeutics in HD.

## Results

### General Animal Health and Inflammation

Consistent with previous reports of *Htt*^*Q111/+*^ mice on the CD1 and B6J strains ^9, 10^, no body weight changes were observed between 3 and 12 months of age (effect of genotype, F_*(1,94)*_ = 3.1, p = 0.07, mean CAG length = 113, cohort summary in Table 1). Important plasma chemistry parameters, including protein, glucose and ion concentrations, as well as liver enzyme levels, were normal at 16-19 weeks of age in *Htt*^*Q111/+*^ mice ^10^. Because we are interested in later-onset phenotypes, we examined a suite of clinical chemistry parameters at 12 months of age - no changes were observed in any of these parameters in male *Htt*^*Q111/+*^ mice at this age (Table 2). Consistent with findings in younger mice, and coupled with body weight observations, these results suggest that up to 12 months of age *Htt*^*Q111/+*^ mice are grossly healthy. Subtle increases in concentrations of pro-inflammatory chemokines and cytokines have been observed in presymptomatic HD mutation carriers ^19^ and some (e.g. YAC128), but not all (e.g. BACHD), transgenic HD mice by 12 months of age ^20^. To understand whether increased peripherally-detectable inflammation is occurring in *Htt*^*Q111/+*^ mice we quantified cytokines, chemokines and acute phase reactants in the plasma of 12-month-old, male *Htt*^*Q111/+*^ mice (multi-analyte profiling, Myriad RBM). We successfully quantified 16 of these molecules (Table 2: Eotaxin, EGF, IP-10, IL-1β, IL-18, LIF, M-CSF-1, MDC, MIP-1α, MIP-3β, MCP-1, MCP-3, MCP-5, Thrombopoietin, TIMP-1, VEGF-A) but observed no changes in 12-month-old *Htt*^*Q111/+*^ mice. Other important pro-inflammatory molecules (FGF9, FGF-basic, GM-CSF, KC/GRO, IFN-γ,IL-1α, IL-2, -3, -5, -6, -7, -10, -11, -12p70, -17a, MIP1-β, OSM, SCF, TNF-α) were also assayed, but fell below the indicated limit of detection in all mice studied (lower limit of quantification, LLoQ, provided in Table 2). These data suggest that peripherally-detectable immune activation is either not occurring in 12-month-old, male *Htt*^*Q111/+*^ mice, or is sufficiently subtle to be missed by these analyses.

**Table 1:** Molecular natural history cohort characteristics.

| Age | Sex | Genotype | N | Measured CAG (SD) | Terminal Body Weight (SD) |
| --- | --- | --- | --- | --- | --- |
| 3 Months | M | B6.Htt<sup>+/+</sup> | 9 |  | 24.3 (1.7) |
| 3 Months | M | B6.Htt<sup>Q111/+</sup> | 9 | 113 (3.3) | 23.8 (1.4) |
| 3 Months | F | B6.Htt<sup>+/+</sup> | 9 |  | 18.2 (0.7) |
| 3 Months | F | B6.Htt<sup>Q111/+</sup> | 9 | 114 (2.5) | 17.2 (1.4) |
| 9 Months | M | B6.Htt<sup>+/+</sup> | 9 |  | 28.2 (1.8) |
| 9 Months | M | B6.Htt<sup>Q111/+</sup> | 9 | 113 (5.0) | 31.4 (1.8) |
| 9 Months | F | B6.Htt<sup>+/+</sup> | 9 |  | 23.3 (1.2) |
| 9 Months | F | B6.Htt<sup>Q111/+</sup> | 9 | 113 (3.3) | 23.5 (1.6) |
| 12 Months | M | B6.Htt<sup>+/+</sup> | 9 |  | 30.6 (2.6) |
| 12 Months | M | B6.Htt<sup>Q111/+</sup> | 9 | ... | 31.4 (1.6) |
| 12 Months | F | B6.Htt<sup>+/+</sup> | 9 |  | 25.2 (3.7) |
| 12 Months | F | B6.Htt<sup>Q111/+</sup> | 9 | ... | 26.1 (2.5) |

**Table 2:**
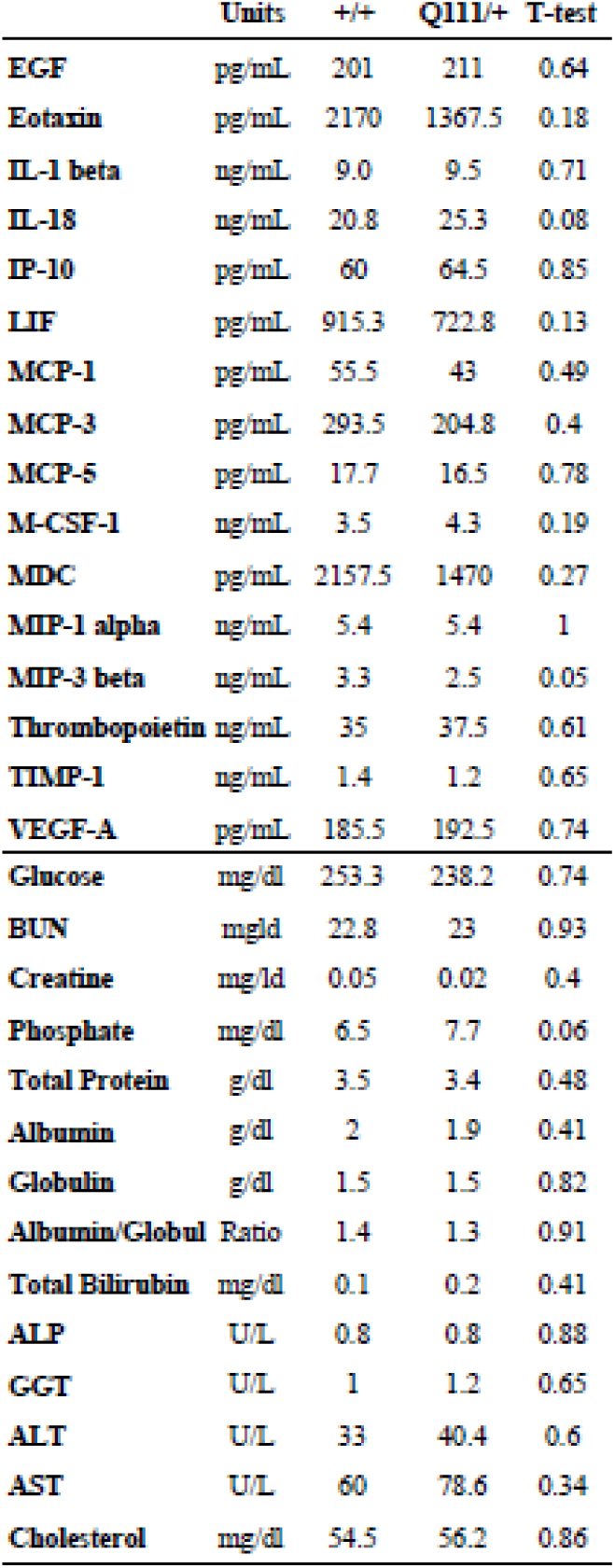
Clinical chemistry parameters and Cytokines. *Clinical chemistry parameters are normal in 12 month old Htt*^*Q111/+*^ *mice. Top level analytes are from inflammationMAP (Myriad RBM, Austin, TX, USA), lower level analytes are from clinical chemistry panel (Phoenix Central Labs, Mukilteo, WA, USA). Abbreviations: epidermal growth factor (EFG), interleukin (IL), interferon gamma-induced protein (IP), monocyte chemoattractant protein (MCP), macrophage colony-stimulating factor (M-CSF), macrophage-derived chemokine (MDC), macrophage inflammatory proteins (MIP), tissue inhibitor of metalloproteinase (TIMP), vascular endothelial growth factor-A (VEGF-A), alkaline phosphatase (ALP), gamma-glutamyl transferase (GGT), alanine aminotransferase (ALT), aspartate aminotransferase (ASP). Inflammation markers measured below LLoQ (LLoQ listed): Fibroblast growth factor (FGF)-9 (2.4 ng/mL), FGF-basic (9.6 ng/mL), granulocyte-macrophage colony-stimulating factor (15 pg/mL), keratinocyte chemoattractant/human growth-regulated oncogene (0.028 ng/mL), interferon gamma (182 pg/mL), IL-1 alpha (100 pg/mL), IL-2 (70 pg/mL), IL-3 (5.9 pg/mL), IL-4 (60 pg/mL), IL-5 (0.6 ng/mL), IL-6 (4.7 pg/mL), IL-7 (0.19 ng/mL), IL-10 (125 pg/mL), IL-11 (56 pg/mL), IL-12p70 (0.091 ng/mL), IL-17A (0.018 ng/mL), MIP-1 beta (142 pg/mL), MIP-2 (11 pg/mL), oncostatin-M (0.24 ng/mL), stem cell factor (652 pg/mL), tumor necrosis factor-alpha (0.082 ng/mL). N = 4 male Htt*^*Q111/+*^, *4 male Htt*^+/+^, *subset of 12-month mice from Table 1)*.

|  | Units | +/+ | Q111/+ | T-test |
| --- | --- | --- | --- | --- |
| EGF | pg/mL | 201 | 211 | 0.64 |
| Eotaxin | pg/mL | 2170 | 1367.5 | 0.18 |
| IL-1 beta | ng/mL | 9.0 | 9.5 | 0.71 |
| IL-18 | ng/mL | 20.8 | 25.3 | 0.08 |
| IP-10 | pg/mL | 60 | 64.5 | 0.85 |
| LIF | pg/mL | 915.3 | 722.8 | 0.13 |
| MCP-1 | pg/mL | 55.5 | 43 | 0.49 |
| MCP-3 | pg/mL | 293.5 | 204.8 | 0.4 |
| MCP-5 | pg/mL | 17.7 | 16.5 | 0.78 |
| M-CSF-1 | ng/mL | 3.5 | 4.3 | 0.19 |
| MDC | pg/mL | 2157.5 | 1470 | 0.27 |
| MIP-1 alpha | ng/mL | 5.4 | 5.4 | 1 |
| MIP-3 beta | ng/mL | 3.3 | 2.5 | 0.05 |
| Thrombopoietin | ng/mL | 35 | 37.5 | 0.61 |
| TIMP-1 | ng/mL | 1.4 | 1.2 | 0.65 |
| VEGF-A | pg/mL | 185.5 | 192.5 | 0.74 |
| Glucose | mg/dl | 253.3 | 238.2 | 0.74 |
| BUN | mg/dl | 22.8 | 23 | 0.93 |
| Creatine | mg/dl | 0.05 | 0.02 | 0.4 |
| Phosphate | mg/dl | 6.5 | 7.7 | 0.06 |
| Total Protein | g/dl | 3.5 | 3.4 | 0.48 |
| Albumin | g/dl | 2 | 1.9 | 0.41 |
| Globulin | g/dl | 1.5 | 1.5 | 0.82 |
| Albumin/Globul | Ratio | 1.4 | 1.3 | 0.91 |
| Total Bilirubin | mg/dl | 0.1 | 0.2 | 0.41 |
| ALP | U/L | 0.8 | 0.8 | 0.88 |
| GGT | U/L | 1 | 1.2 | 0.65 |
| ALT | U/L | 33 | 40.4 | 0.6 |
| AST | U/L | 60 | 78.6 | 0.34 |
| Cholesterol | mg/dl | 54.5 | 56.2 | 0.86 |

### Striatal Transcriptional Alterations

Altered transcription is an early and prominent feature of HD pathology, and is observed in both animal models and samples from human patients ^12^. Because we are interested in intervening at the earliest possible point in the pathogenic process, we investigated the time course of transcriptional changes in the central nervous system of *Htt*^*Q111/+*^ mice. As an initial, untargeted pilot study, we examined striatal and cerebellar genome-wide transcriptional changes in 3- and 9-month-old *Htt*^*Q111/+*^ mice using mRNA sequencing (RNASeq). At 3 months of age, no robust transcriptional alterations were detected in the striatum of *Htt*^*Q111/+*^ mice compared to wild-type mice (Figure 1a, 5 quantified transcripts have an effect of genotype with a false discovery rate (FDR) of < 5%). By 9 months of age, 726 transcripts were changed in the striatum of *Htt*^*Q111/+*^ mice compared to wild-type mice (Figure 1a, 481 down-regulated, 245 up-regulated at FDR < 5%). The cerebellum, despite robustly expressing mutant *Htt* ^21,22,23^, is relatively spared from pathological tissue loss in human HD patients ^24^ and the YAC128 mouse model of HD ^14^. We therefore compared the rate and scale of transcriptional dysregulation between the cerebellum and striatum of *Htt*^*Q111/+*^ mice at 3 and 9 months of age. There were very few significant effects of CAG expansion in Htt in the cerebellum at either age (Figure 1a, 3 genotype-sensitive transcripts, 2 down-regulated, 1 up-regulated at FDR < 5%). These data confirm that, as in humans with HD and other animal models, progressive striatal transcriptional dysregulation is a feature of aging *Htt*^*Q111/+*^ mice.

We next examined pathway enrichment of dysregulated transcripts using both hypergeometric ^25^ and gene set enrichment analysis (GSEA) ^26^ techniques. The hypergeometric test examines over-enrichment of differentially expressed genes in a specific pathway, while GSEA quantifies whether a set of pathway genes is over-represented at the top or bottom of an ordered list of genes - here, the full list of 9-month striatal transcripts, ordered by the fold-change in *Htt*^*Q111/+*^ mice compared to wild-type mice. Hypergeometric testing of 726 genotype-sensitive (FDR < 5%) striatal transcripts (from a gene universe of 19,031 robustly assayed genes) revealed significant enrichment of striatal genotype-sensitive transcripts in 8 pathways with an adjusted p-value < 0.05, notably in neuronal signaling pathways (Figure 1b). GSEA revealed 15 gene sets significantly enriched at the high or low end of the list of striatal transcripts, ordered by their genotype expression ratio (*Htt*^*Q111/+*^ / *Htt^+/+^*; Figure 1c). Nineteen common pathways had *nominal* enrichment p-values < 0.05 in both GSEA and hypergeometric analyses, including Reactome pathways involved in neurotransmission and cellular signaling (e.g. transmission across chemical synapses, neuronal system, signaling by GPCR, GPCR downstream signaling; Table S1), suggesting synaptic and signaling transcripts are altered at the transcriptional level in the 9-month-old *Htt*^*Q111/+*^ striatum.

**Figure 1.**
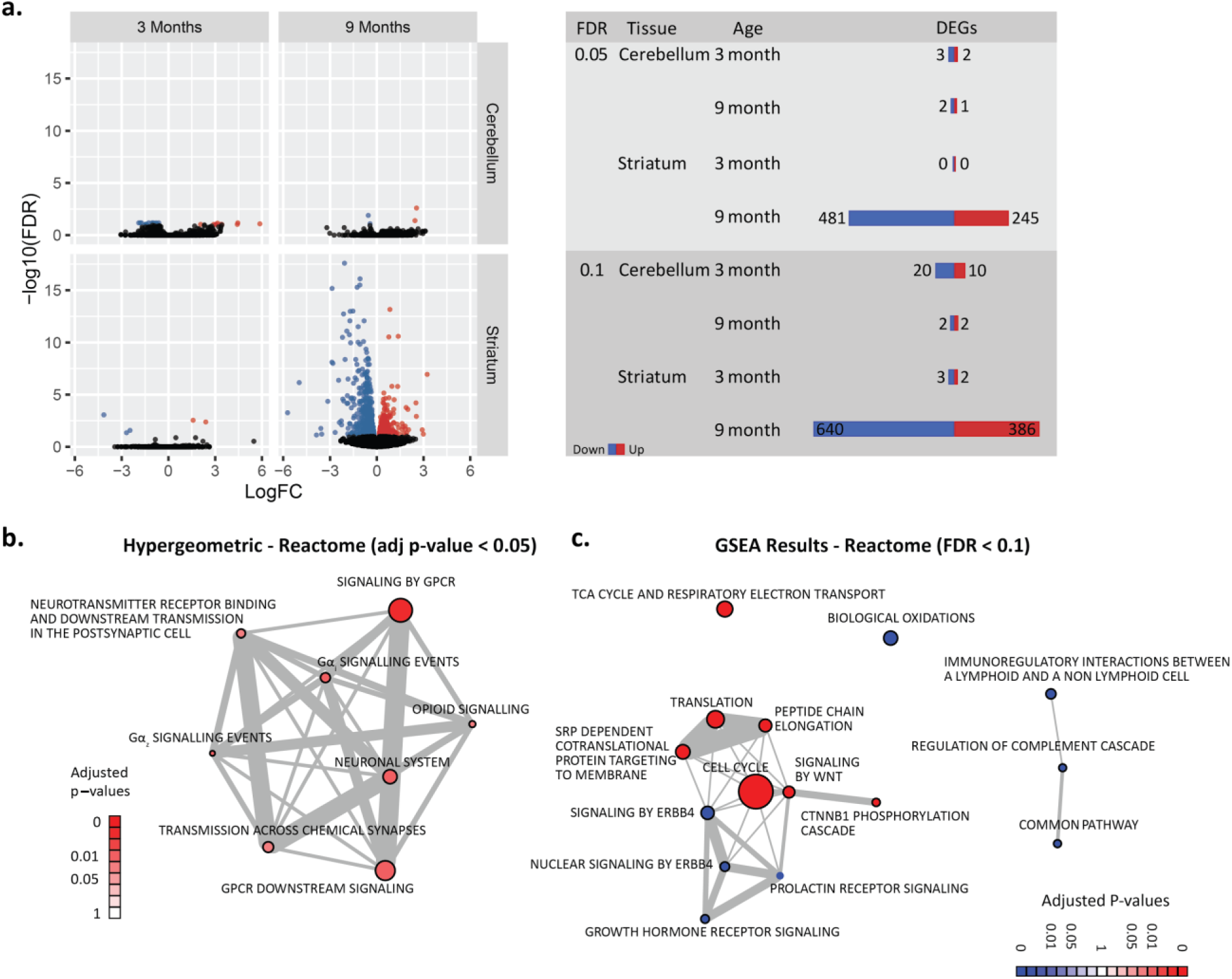
*Progressive, striatal-specific, transcriptional dysregulation in the aging Htt*^*Q111/+*^ *brain.* ***a.** The impact of genotype on the expression level of each of the 19,031 transcripts assayed is indicated - statistical significance on the y-axis (−log_10_ FDR), and fold-change on the x-axis (log fold-change, Htt*^*Q111/+*^ / *Htt*^+/+^*). Color indicates statistical significance - black indicates FDR > 0.1, red and blue indicate FDR < 0.1 for up- and down-regulated genes in the Htt*^*Q111/+*^ *striatum, respectively. **b.** Network diagram depicting the 8 reactome pathways with adjusted p-values less than 0.05 in a hypergeometric analysis, and the proportion of genes shared between each pathway. The size of each node corresponds to the overall gene set size (e.g. signaling by GPCR pathway includes 853 genes, Gα_z_ signalling events 44), while the width of the edges corresponds to the Jaccard similarity coefficient for the pair of gene sets (the intersection of the two sets divided by its union). **c.** Network diagram depicting the 15 reactome pathways whose genes are non-randomly distributed on the ordered list of striatal transcripts with an FDR < 0.1, and the proportion of genes shared between each pathway. Blue nodes indicate down-regulated pathways, while red nodes indicate upregulation in the 9 month old Htt*^*Q111/+*^ *striatum.*

To orthogonally validate the observed striatal transcriptional alterations and examine their relevance as useful preclinical tools in *Htt*^*Q111/+*^, we quantified specific transcript levels in the striatum of 3-, 9- and 12-month-old *Htt*^*Q111/+*^ mice using quantitative real-time polymerase chain reaction (QRT-PCR). As predicted by cross-sectional RNASeq data (Figure 1), transcript levels of the critical striatal signaling gene dopamine- and cAMP-regulated phosphoprotein, Mr 32 kDa (DARPP32) are normal in *Htt*^*Q111/+*^ mice at 3 months of age, but reduced at both 9 and 12 months of age (effect of genotype F_(1,47)_=6.74, p = 0.01, Figure 2a). Similarly, we find that levels of a suite of additional neuronal signaling and synaptic genes (including: Cnr1, Drd2, Homer1, Pde10a, Penk, Scn4b) are reduced in aged (9- and 12-month-old) but not young (3-month-old) *Htt*^*Q111/+*^ mice (Figure 2a, effect sizes for each target shown in Figure 2b). Other transcripts increase over time in *Htt*^*Q111/+*^ and include genes involved in DNA damage (e.g. N4bp2) or immune response (e.g. Islr2, H60b, Figure 2a). These results confirm the findings of our RNAseq study, and suggest that levels of a range of transcripts are sensitive genotype markers in the aging *Htt*^*Q111/+*^ striatum.

**Figure 2.**
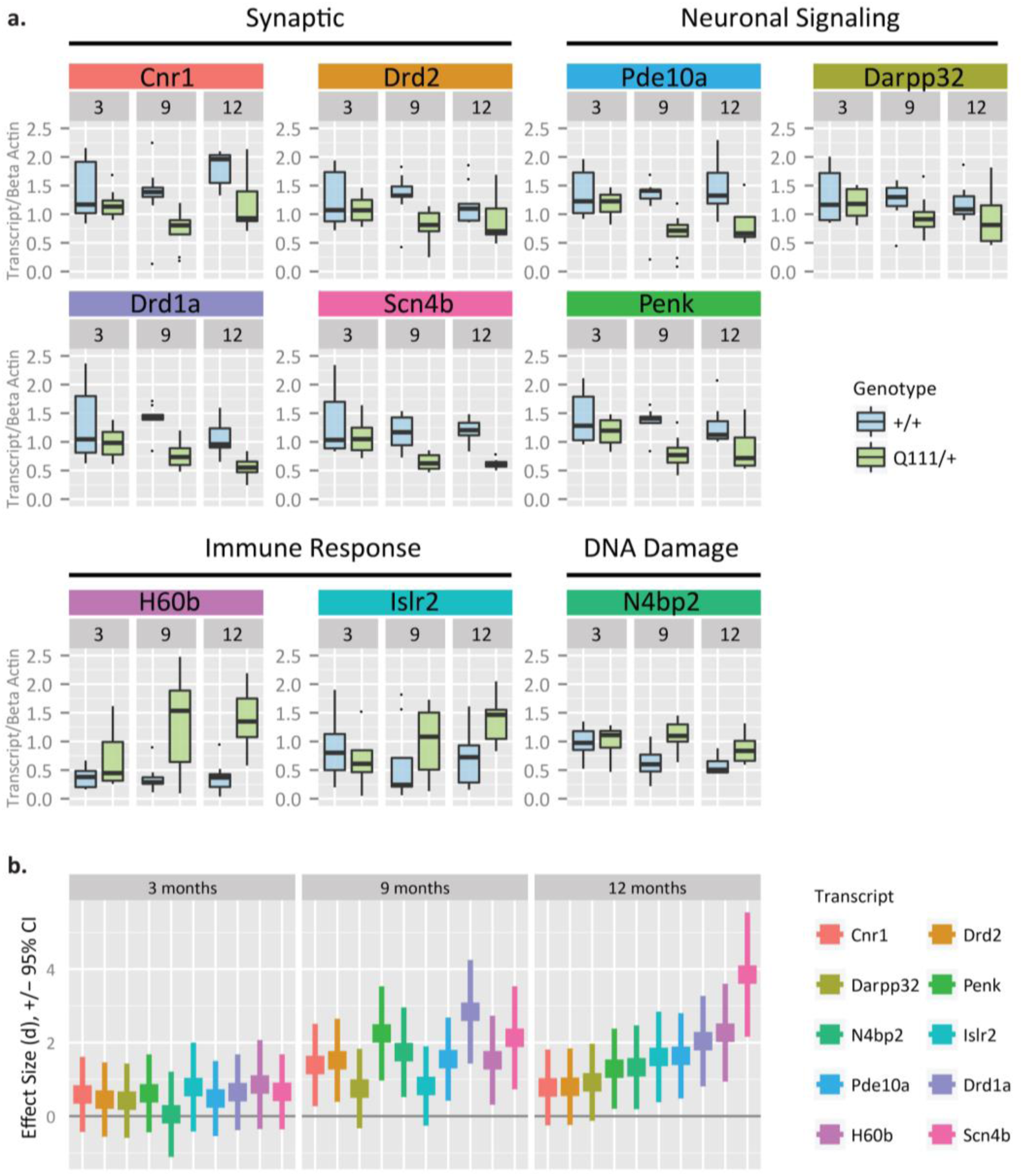
*Confirmation of progressive transcriptional dysregulation in the striatum of aging Htt*^*Q111/+*^ *mice using quantitative real-time polymerase chain reaction.* ***a**. Guided by transcriptional discovery with RNAseq, we quantified a number of transcripts and found that many synaptic and neuronal signalling transcripts were down as mice aged from 3 months to 9- and 12-months. As anticipated, we also found upregulated transcripts related to immune and DNA damage pathways (N = 60, subset of 5 mice per row from Table 1). **b**. Corresponding longitudinal effect sizes for each transcript (highlighted in colored bands above) are displayed in the same color below, with whiskers representing the 95% confidence interval range. Along the x-axis, transcripts are ordered by increasing effect size at 12-months, though robust effects are seen at both 9- and 12-months. Our results demonstrate that these transcripts are sensitive genotype markers that make suitable targets for assessing rescue in interventional trials using the Htt^*Q111/+*^ mouse model.*

### Striatal Histology

We also considered a range of histological endpoints in the dorsolateral striatum of 3-, 9- and 12-month-old *Htt*^*Q111/+*^ mice. Compared to tissue-level analyses such as RNASeq and QRT-PCR, immunohistochemistry (IHC) enables the identification of specific cell types and analysis of subcellular localization. First, a trivial explanation for the observed transcriptional alterations in the striatum of the *Htt*^*Q111/+*^ mice is that the cellular composition or relative cell sizes within the striatum has changed. To examine this possibility, we counted putative neurons, astrocytes and microglia in the dorsolateral striatum (respectively: NeuN-, glial fibrillary acidic protein- and allograft inflammatory factor 1-immunoreactive cells). Between 3-12 months of age, the neuronal density in the dorsolateral striatum of *Htt*^*Q111/+*^ mice does not change (Figure 3a, F_(1, 100)_= 0.03, p = 0.87), nor does the cross-sectional area of NeuN-immunoreactive soma in the dorsolateral striatum at 9 months (NeuN-immunoreactive cell size distributions shown in Figure 3b, two-sample Kolmogorov-Smirnov test D = 0.02, p = 0.9). We next quantified astrocytic and microglial density in the dorsolateral striatum of 12 month old *Htt*^*Q111/+*^ and *Htt*^+/+^ mice, finding that neither is increased in 12 month old *Htt*^*Q111/+*^ mice (Fig 3d). Progressive loss of synaptophysin density has been reported in transgenic HD and other neurodegenerative disease mouse models ^27^. We quantified synaptophysin density in the dorsolateral striatum and find that *Htt*^*Q111/+*^ and *Htt*^+/+^ mice have equivalent levels of synaptophysin staining from 3-12 months of age (Figure 3c, F_(1, 93)_= 3.71, p = 0.057). These results suggest that the robust transcriptional alterations we observe in the aging *Htt*^*Q111/+*^ striatum (Figure 1) are not due to altered numbers or size of neurons, loss of synaptic compartments and that overt gliosis is not a component of the phase of HD modeled by the *Htt*^*Q111/+*^ mice.

**Figure 3.**
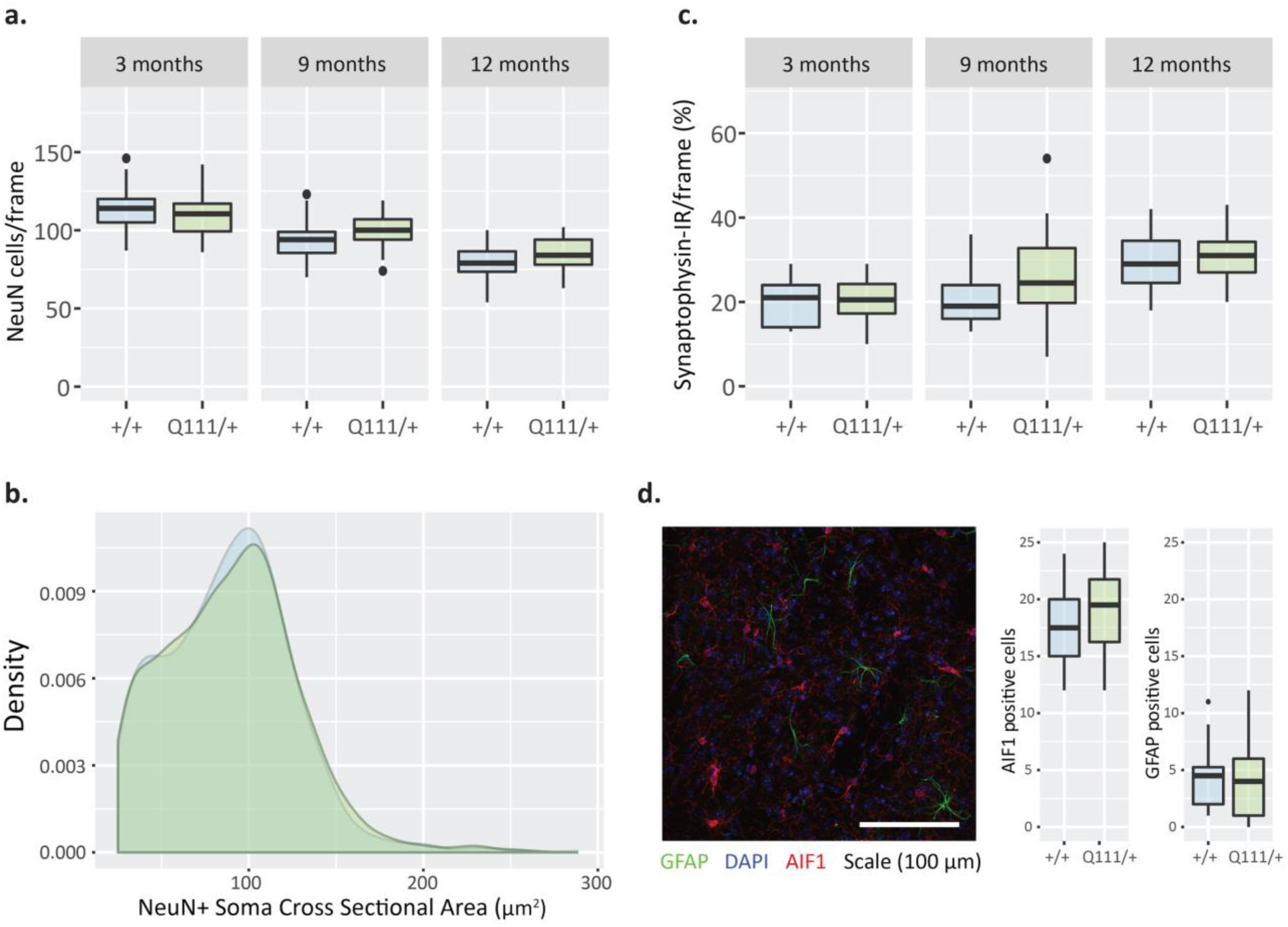
*No evidence of changes in striatal cell distribution, neuronal cell size or a synaptic marker in Htt*^*Q111/+*^ *mice through 12 months of age.* ***a.** NeuN+ cell density is not changed between Htt^*Q111/+*^ and Htt*^+/+^ *mice at 3, 9, or 12 months of age (N = 108, breakdown Table 1). **b.** Distribution of striatal NeuN+ cell sizes does not differ between Htt^*Q111/+*^ and Htt^+/+^ striata at 9 months of age (N = 108, breakdown Table 1). **c.** Synaptophysin staining intensity was examined at 3, 9, and 12 months of age, showing no differences between Htt^*Q111/+*^ and Htt^+/+^ mice. (N = 108, breakdown Table 1). **d.** Glia counts at 12 months show no difference in numbers of GFAP+ astrocytes (green) and Aif1+ microglia (red) in Htt^*Q111/+*^ versus Htt^+/+^ striata. Representative Htt^+/+^ section shown (N = 36, 12-month cohort from Table 1).*

We also examined the appearance of aggregated striatal huntingtin using the MW8 antibody, which detects aggregated, but not diffuse, mutant huntingtin ^28^, but to our knowledge has not yet been applied to study *Htt*^*Q111/+*^ mice. We restricted our analyses to neuronal cell bodies, immunoreactive for NeuN, a well-described pan-neuronal marker ^29^. In the dorsolateral striatum of 3-month-old *Htt*^*Q111/+*^ mice there is virtually no neuronal MW8 immunoreactivity, but by 9 months of age we observe robust accumulation of both small punctate and large inclusion staining in neuronal nuclei (Figure 4a). Between 9 and 12 months of age, increased levels of neuronal intranuclear inclusions (NII) in the *Htt*^*Q111/+*^ striatum are accompanied by a reduction in the total nuclear MW8 immunoreactivity. Total neuronal nuclear MW8 immunoreactivity increases from 3 to 9 months of age, before declining at 12-months of age (genotype F_(1,95)_= 25.92, p < 0.0001, genotype/age interaction F_(2,95)_=10.4, p < 0.0001, Figure 4b). This rise and fall in total neuronal nuclear MW8 immunoreactivity is accompanied by a progressive increase in the percentage of cells with large NIIs, from 0% at 3-months to 13% at 9-months and 28% by 12 months of age (genotype F_(1,99)_=1 56.9, p < 0.0001, genotype/age interaction F_(2,99)_ = 61.5, p < 0.0001, Figure 4d). In parallel, the average size of striatal NIIs in *Htt*^*Q111/+*^ mice increases more than 40% from 1.3 ± 0.7 μm at 9 months to 1.8 ± 0.9 μm at 12 months of age (two-sample Kolmogorov-Smirnov test, D = 0.29, p < 0.0001). Impaired autophagy has been proposed to contribute to impaired proteostasis in HD ^29,30^, and in the brain mHTT interacts directly with p62/Sqstm1, an autophagic receptor protein important for selective macroautophagy ^31^. We observed complete co-localization between p62- and MW8 immunoreactivity in striatal NII’s in *Htt*^*Q111/+*^ mice (Figure 4c) suggesting that, as has been observed in cell culture ^31^, p62 is found in NIIs. We finally considered whether quantitative IHC for specific MSN targets is superior to QRT-PCR quantification. As a proof of concept, we quantified neuronal DARPP32 levels in corticostriatal sections. Consistent with mRNA reductions (Figure 2), we observe reduced neuronal somatic DARPP32 levels in the *Htt*^*Q111/+*^ striatum (Figure 5; genotype F_(1,98)_= 6.81, p = 0.01). We find that, for DARPP32, quantification of IHC data is more variable than QRT-PCR when establishing reductions in the *Htt*^*Q111/+*^ striatum.

**Figure 4.**
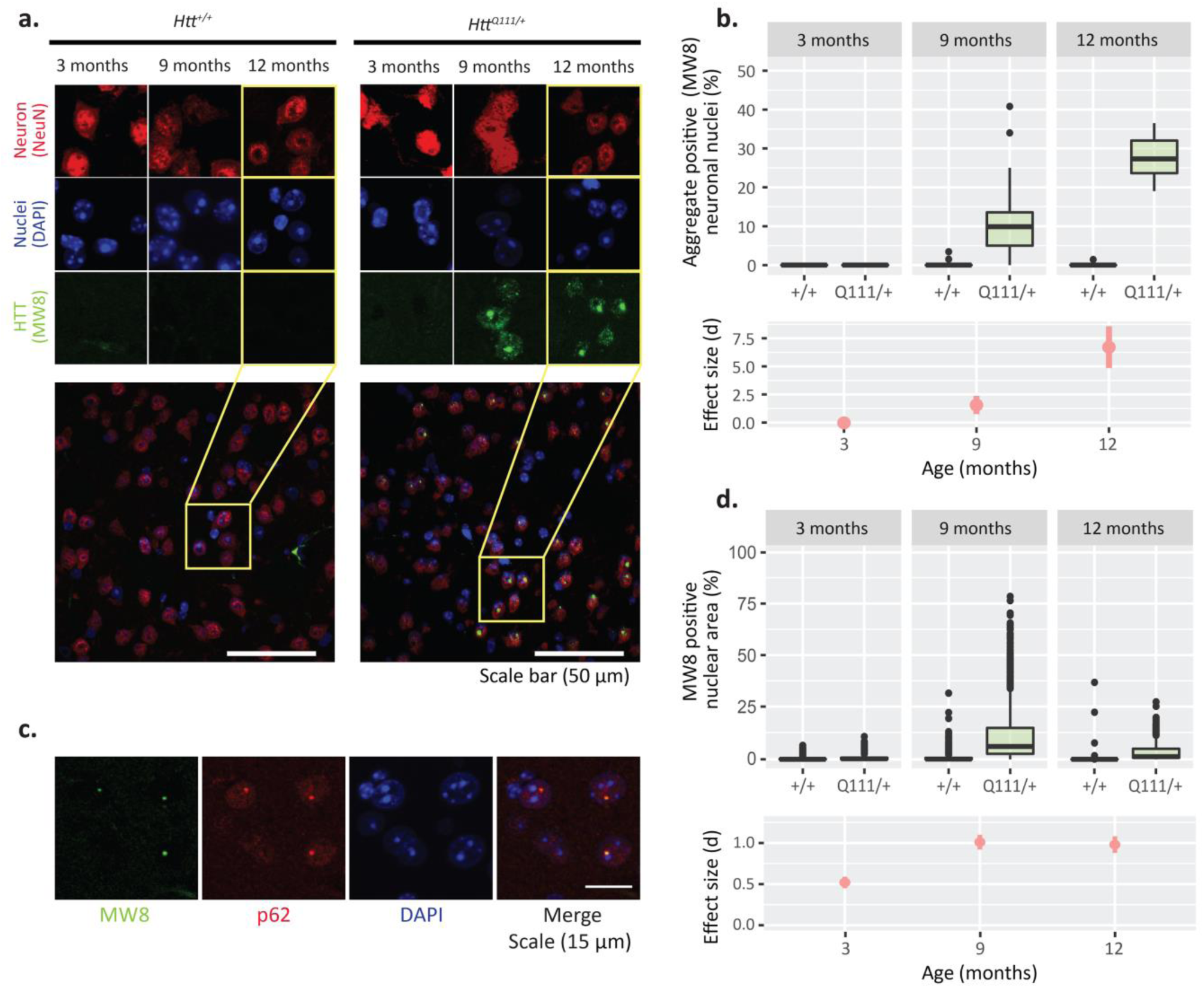
Progressive accumulation of p62+− and huntingtin-immunoreactive neuronal intranuclear inclusions (NIIs) in the striatum of aging Htt^Q111/+^ mice. **a**. Images of the dorsolateral striatum taken in 5-μm sections of 12-month mice triple labelled for neurons (Red; NeuN), nuclei (Blue; DAPI) and aggregated huntingtin (Green; MW8). Accumulation of both large neuronal-nuclear HTT aggregates, as well as small huntingtin nuclear speckles is present in Htt^Q111/+^, but absent in Htt^+/+^ mice. **b**. Using a neuronal (NeuN) mask with particle inclusion from 0.5-5 μm^2^ shows a median of 10% of neurons contain large nuclear aggregates by 9-months, which increases to 27% at 12 months of age. The presence of NIIs provides a robust measure for huntingtin accumulation with an extremely robust effect size for genotype comparisons at 9- and 12-months (d = 1.6 and 6.7, respectively). ANOVA: genotype F_(1,94)_ = 156.91, p < 0.0001, age F_(2,94)_ = 63.87, p < 0.0001; genotype x age F_(2,94)_ = 61.48, p < .0001; N = 108, breakdown Table 1. **c**. Images co-labelled for aggregated huntingtin (Green; MW8), and autophagy adaptor protein p62 (Red), demonstrate p62 is colocalized with aggregated huntingtin in 12-month Htt^Q111/+^ mice. **d.** As with MW8, measuring nuclear immunoreactivity of p62 provides a robust measure of large nuclear aggregates; ANOVA: genotype F_(1,99)_ = 40.22, p < 0.0001, age F_(2,99)_ = 22.82, p < 0.0001; genotype x age F_(2,99)_ = 14.25, p < .0001; N = 108, breakdown in Table 1.

**Figure 5.**
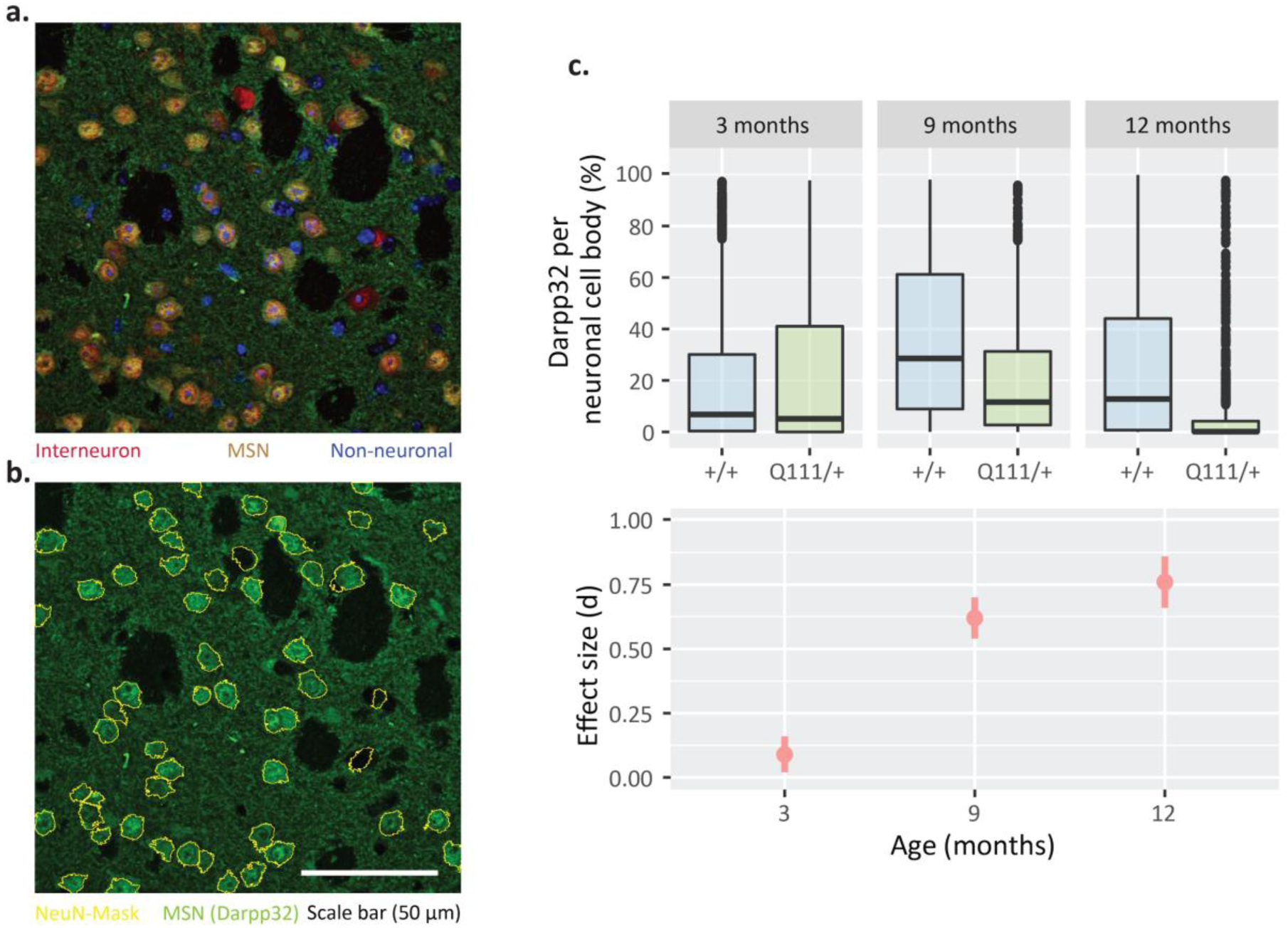
*Histological analysis confirms protein-level reductions in DARPP32 in the aging Htt*^*Q111/+*^ *striatum*. ***a.** Representative image of DARPP32, NeuN, and DAPI triple-labelled dorsolateral striatum highlights interneurons (Red), medium spiny neurons (MSN; Yellow), and non-neuronal cells (Blue). **b.** DARPP32 staining with NeuN positive cells (Yellow) demonstrate the inclusion areas for DARPP32 quantification. **c.** DARPP32 immunofluorescence per neuronal cell body is reduced in* Htt^Q111/+^ *mice (ANOVA: Genotype F_*(1,98)*_ = 6.81, p = 0.01, N = 108, breakdown in Table 1).*

### Behavioral Analyses

We cross-sectionally analyzed behavior in several cohorts of 3-9 month old *Htt*^*Q111/+*^ mice, including assays of motivation, anhedonia, depression, and anxiety (cohorts described in methods). We first measured reward seeking behavior in *Htt*^*Q111/+*^ mice using an operant fixed ratio 1 (FR1) behavioral assay to examine how many sweet rewards mice will perform for in a 5 minute block. We found that *Htt*^*Q111/+*^ mice demonstrate reduced reward attainment relative to *Htt*^+/+^ mice (Figure 6a; Genotype: *F*_(1,14)_ = 17.0, *p* = 0.001: Genotype x Reward Size: *F*_(2,28)_ = 8.4, *p* = 0.001). In an attempt to discern hedonic from motivational explanations for this reduced reward attainment, we employed a sucrose pellet consumption task, and found that overall sucrose preference increased in the *Htt*^*Q111/+*^ mice compared to *Htt*^+/+^ mice (Figure 6e; Genotype:*F*_(1,14)_ = 15.9, *p* = 0.001). In a separate cohort of mice, we also examined preference for either 2% or 4% sucrose in drinking water, and found no difference between 9 month old *Htt*^+/+^ and *Htt*^*Q111/+*^ mice (2% sucrose Genotype x Sucrose concentration: *F*_(1,38)_ = 0.03, *p* = 0.87; 4% sucrose Genotype x Sucrose concentration: *F*_(1,35)_ = 0.41, *p* = 0.53, Figure S1). To explore the potential contributions of anxiety or depression to the observed reward attainment changes we conducted several tests of these phenotypes in a separate cohort of 9-month old *Htt*^+/+^ and *Htt*^*Q111/+*^ mice. First we conducted the Porsolt swim test, a measure of behavioral despair ^30^, where we observe no differences between 9-month old *Htt*^+/+^ and *Htt*^*Q111/+*^ mice in the duration of time spent inactive during the task (Figure 6f; *t*(38) = 0.8, *p* = 0.9). Similarly, using the elevated plus maze task as a measure of anxiety ^32^, we found that 9-month old *Htt*^+/+^ and *Htt*^*Q111/+*^ mice do not differ in the amount of time spent in the open arms (Figure 6g; *t*(37) = 0.7, *p* = 0.5), or in the percentage of total arm entries into the open arms, (*t*(37) = -1.3, *p* = 0.2). As a final measure of anxiety levels in *Htt*^*Q111/+*^ mice, we also employed the light/dark exploration task ^32,33^, finding that *Htt*^+/+^ and *Htt*^*Q111/+*^ mice spend the same amount of time in the light and dark halves of the apparatus (Figure 6h; *t*(38) = 0.04, *p* = 0.9) and enter the light compartment of the maze a similar number of times (*t*(38) = −0.6, *p* = 0.5). Altogether, these data suggest that up to 9-months of age *Htt*^*Q111/+*^ mice do not display anxiety-related or depressive-like symptoms, and declines observed in reward seeking behavior may reflect presently undefined cognitive or motivational issues worthy of additional study.

**Figure 6.**
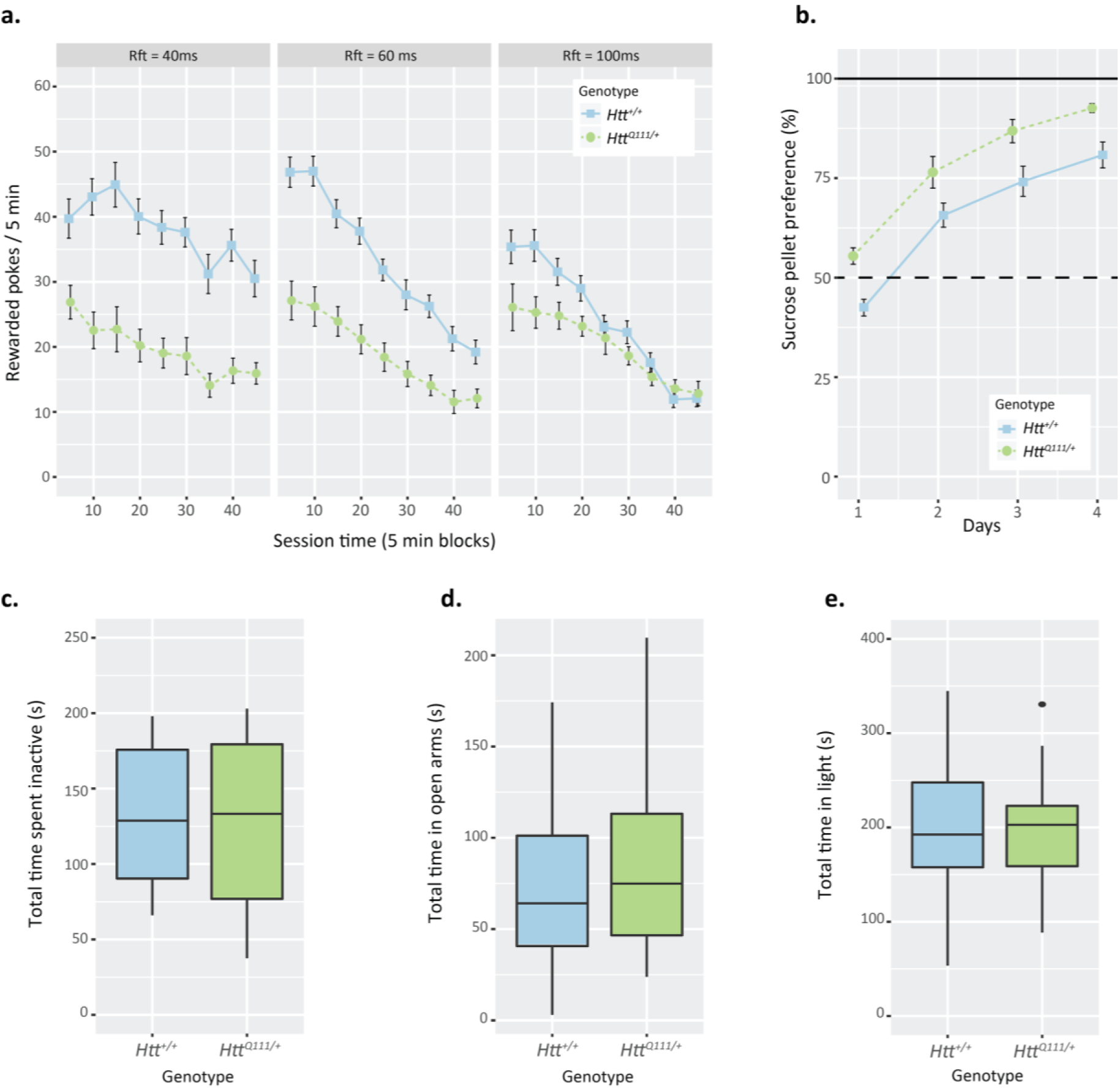
*Presence of Htt*^*Q111*^ *allele results in reduced reward seeking behavior independent of anxiety-like or depressive behavior.* ***a.** Htt^*Q111/+*^ mice display reduced reward attainment compared to Htt*^+/+^ *mice* across three reinforcement durations (Rft = 40 ms, 60 ms, & 100 ms with 100 ms being 5µl) *during FR1 operant testing (Genotype: F*_*(1,14)*_ *= 17.0, p = 0.001, Genotype x Reward: F*_*(2,28)*_ *= 8.4, p = 0.001). **b.** Htt*^*Q111/+*^ *mice exhibit a stronger preference for sucrose pellets during the hours sucrose consumption task compared to Htt*^+/+^ *mice (Genotype: F*_*(1,14)*_ *= 15.9, p = 0.001). **c.** Htt*^*Q111/+*^ *and Htt*^+/+^ *mice do not differ in the amount of time spent inactive during the Porsolt swim task. **d.** Htt*^*Q111/+*^ *and Htt*^+/+^ *mice spend a similar amount of time in the open arms of the elevated plus maze. **e.** Htt*^*Q111/+*^ *and Htt*^+/+^ *mice do not differ in the amount of time spent in the light compartment of the apparatus during the light/dark exploration task.*

### Power Analysis for preclinical studies in Htt^Q111/+^ mice

To establish the utility of our natural history data, we conducted several power analysis studies to understand whether these results would be useful in a preclinical trial setting. We considered a hypothetical 2 × 2 factorial experiment, with two genotypes (*Htt*^+/+^ vs. *Htt*^*Q111/+*^) and two treatment groups (baseline vs. a hypothetical treatment). We simulated data for *Htt*^+/+^ and *Htt*^*Q111/+*^ mice in the ‘treatment’ condition by drawing values from a random normal distribution with the same mean, variance, and covariance as our real natural history data. Assuming a 50% rescue from a hypothetical intervention, we established the power of single molecular endpoints in a preclinical study with 10 animals per arm (4 arms - *Htt*^+/+^ and *Htt*^*Q111/+*^ mice in a treatment or control arm). By 12 months of age, several individual endpoints provide reasonable power to detect this 50% rescue, including reductions in striatal *Scn4b* mRNA levels and an increase in MW8-immunoreactive aggregate counts (Figure 7a). We next considered whether combining information from multiple molecular endpoints would improve the power to detect a partial rescue. We trained an elastic net logistic regression model to classify *Htt*^*Q111/+*^ versus *Htt*^+/+^ mice using a weighted combination of the QRT-PCR and IHC endpoints in the striatum. This model distinguished *Htt*^*Q111/+*^ vs. *Htt*^+/+^ mice with >90% accuracy in training data from 9- and 12-month-old mice. The model assigns non-zero weights to 0 endpoints in 3-month-old mice, 9 in 9-month-old mice, and 9 in 12-month-old mice (regression coefficients for individual endpoints provided in Figure 7d). With a sample size of 10, this multivariate model would have 80% power to detect a 35% rescue in 12-month-old mice or a 60% rescue in 9-month-old mice (Figure 7b). With n=20, the model had 80% power to detect a 25% rescue in 12-month-old mice or a 40% rescue in 9-month-old mice (Figure 7c). These results suggest that combining information from multiple molecular endpoints can improve power to detect subtle effects of treatments.

**Figure 7.**
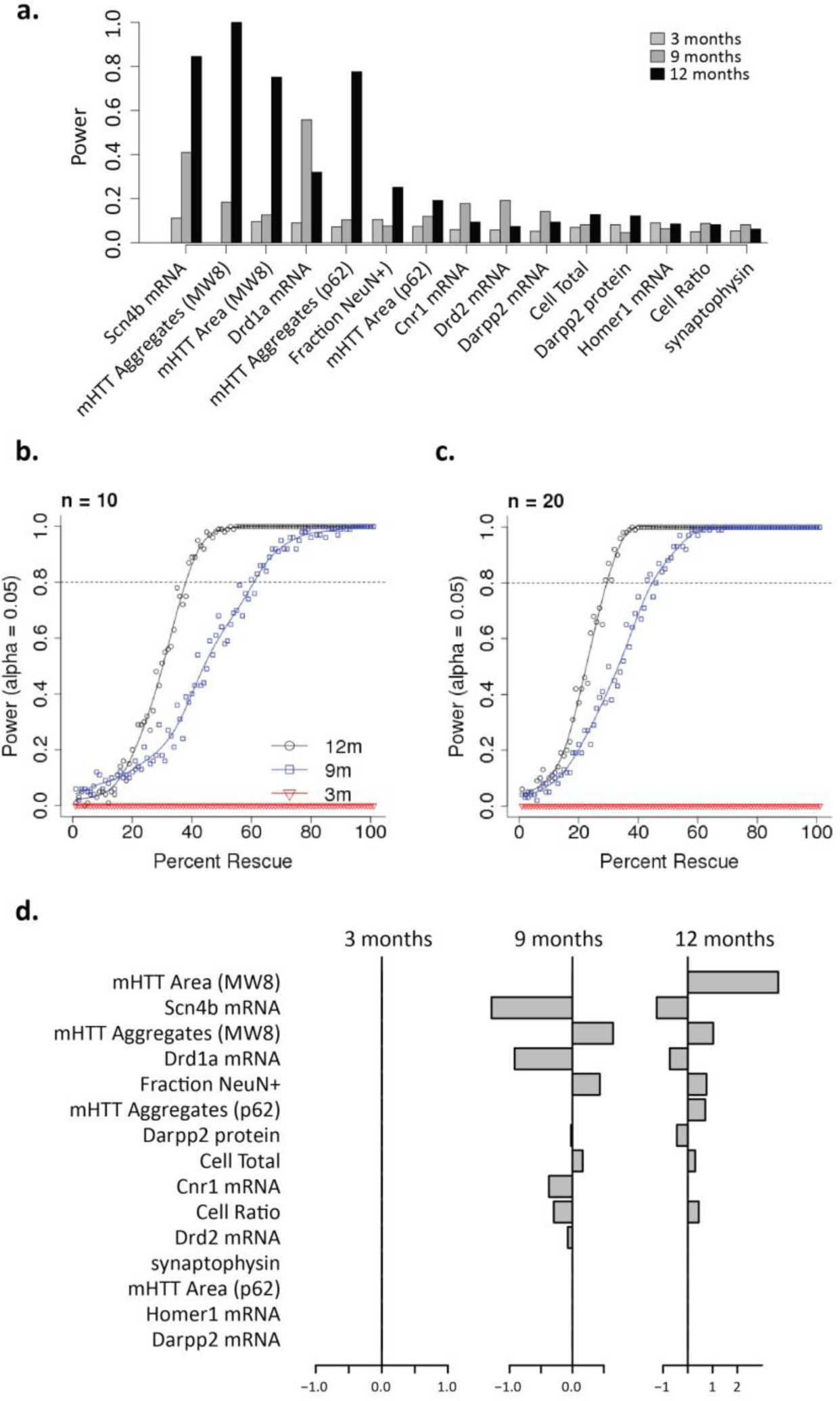
Power analysis suggests multivariate endpoints maximize power for pre-clinical study. ***a.** Estimated power to detect a 50% rescue of individual endpoints in 3-, 9- or 12-month-old mice, assuming a sample size of 10 Htt*^*Q111+*^ *mice and 10 Htt*^+/+^ *mice in baseline conditions and 10 Htt*^*Q111+*^ *mice and 10 Htt*^+/+^ *mice given a hypothetical treatment that induces a partial rescue. **b,c.** Power to detect partial rescue at each timepoint using a multivariate elastic net classifier and a sample size of 10 **(b)** or 20 **(c)** mice per group. **d.** Regression coefficients (weights) assigned to each biomarker in the elastic net model. Positive and negative coefficients indicate that the marker is increased or decreased in Htt*^*Q111/+*^ *mice, respectively.*

## Discussion

We have here characterized the health and progression of phenotypes in the *Htt*^*Q111/+*^ mouse, which accurately model the zygosity and *Htt* expression level of human HD patients. We found that, in contrast to transgenic N-terminal HTT fragment models of the disease, which have rapidly progressing and terminal disease, the *Htt*^*Q111/+*^ mice were grossly healthy through 12 months of age in terms of body weight, plasma chemistry and both central and peripheral inflammation, extending previous observations of these mice ^10^. Despite their generally healthy state in their first year of age, during this interval they presented striatal-specific, progressive, molecular changes consistent with those observed in other animal models of HD and indeed human mutation carriers, as well as specific behavioral phenotypes consistent with early HD. Using simulation studies, we demonstrated that these animals provide a powerful tool for experimental therapeutics targeting these early molecular changes.

In a recent review, Menalled and Brunner ^11^ demonstrated that less than 5% of preclinical studies of reviewed therapeutic agents in HD were tested in knock-in models of the disease. The R6 and N171 lines of mice, which transgenically express short fragments of mHtt, were much more commonly used, together comprising 72% of all preclinical studies examined. Translation from animal models of neurodegeneration to human clinical trials has been disappointing, with a large number of agents predicted to be useful in transgenic animals failing to provide clinical benefit in human HD, for example ^11^. A major benefit of screening therapeutics in short fragment mouse models of HD is that they present robust behavioral phenotypes, including motor alterations, affective impairments and cognitive alterations ^4^. Similar, though less pronounced, phenotypes have been observed in full-length YAC128 and BACHD mice ^9^. Distinguishing central effects from peripheral ones is difficult in these animals because they either progressively lose (e.g. R6/2 ^34^ and N171 ^34,35^) or gain (YAC128 ^36^ and BACHD ^9^) significant amounts of body weight in parallel with the development of progressive molecular and behavioral phenotypes. Large observational studies have suggested that HD patients may, on average, lose weight ^9,37^, as observed in short fragment mouse models of HD. However, more recently, careful and well-powered studies of presymptomatic HD mutation carriers and early stage patients reveals they have normal body composition and plasma chemistry ^38^, consistent with our observations in aging *Htt*^*Q111/+*^ mice (Table 1).

While presymptomatic HD mutation carriers do not show clinically obvious motor impairments, by definition, they do show clear sub-clinical cognitive impairments ^39^ and progressive affective alterations. Amongst affective changes in HD mutation carriers, apathy has emerged as the most widely observed ^40^, and uniquely amongst neuropsychiatric symptoms, progresses in a continuous fashion with disease state ^40,41^. Like presymptomatic HD mutation carriers, previous studies of *Htt*^*Q111/+*^ mice revealed normal gross motor behavior and grip strength through 12 months of age ^9,10,42^. Recent work with q175 HD knock-in mice suggests they present subtle motivational alterations detected using mixed fixed-/progressive-ratio operant tasks ^43^. Similarly, reduced executive function has been observed in *Htt*^*Q111/+*^ mice using delayed nonmatch to sample tasks ^42^. The data presented here (Figure 6) suggest that 9 month old *Htt*^*Q111/+*^ mice do not have enhanced anxiety or depression-like behaviors, but do show motivational deficits, most notably a reduced performance on a fixed-ratio 1 task, despite normal hedonic drive for sweet stimuli at this time point. These behavioral studies, in conjunction with the body weight and general health data presented here, suggest that *Htt*^*Q111/+*^ mice more closely resemble pre-symptomatic mutation carriers than do short fragment models of HD. Importantly, this suggests interventions targeting apathy, the single most prevalent and progressive psychiatric manifestation of HD ^41, 40^, can be tested in knock-in models of HD.

Transcriptional dysregulation in HD has long been noted in both human samples and samples taken from experimental models. Here, using mRNA sequencing techniques we provide quantitative evidence about the degree to which striatal transcriptional changes exceed transcriptional changes in a relatively spared tissue in HD, the cerebellum. Indeed, at 9 months of age, the striatum has more than 250-fold more differentially expressed transcripts than does the cerebellum at this age (Figure 1A). This is consistent with the fact that the cerebellum is grossly spared from pathological volume change in both human patients ^24^ and full-length mouse models of HD ^14^. A concern with transcriptional changes in pathologically affected tissue is that tissue architecture changes (in either cellular composition or relative cell size changes) may explain alterations in mRNA abundance in tissue level analyses. The evidence presented here suggests that robust transcriptional dysregulation in the striatum of *Htt*^*Q111/+*^ mice precedes any alterations in the density of striatal neurons, astrocytes, microglia or synapses. In fact, of the presented phenotypes, altered transcript levels are amongst the most powerful features for distinguishing *Htt*^+/+^ from *Htt*^*Q111/+*^ mice at 9-12 months of age (Figure 7). These data confirm that striatal transcriptional dysregulation occurs in the absence of changes in cell number in early HD, occurring the absence of neurodegeneration, gliosis or overt loss of synapses. Also consistent with previous observations of *Htt*^*Q111/+*^ mice at younger ages ^17^, we observe progressive striatal-specific accumulation of neuronal intranuclear inclusions (NIIs) in *Htt*^*Q111/+*^ mice. Similar to aggregates found in human HD, these large inclusions are primarily restricted to neurons (we do not observe any glial NIIs), and co-stain with autophagic cargo marker p62 (Figure 4). Our power analyses suggest that treatments that alter either mHTT levels, or improve proteostasis and thereby alter aggregate formation, can be robustly tested *in vivo* using the *Htt*^*Q111/+*^ mice with adequate power.

In summary, we present data suggesting that the *Htt*^*Q111/+*^ mice are grossly healthy during their first year of life, yet present a range of molecular and behavioral alterations consistent with presymptomatic HD mutation carriers. Our power analyses revealed that these changes can provide sufficiently powered endpoints for preclinical studies of neuroprotective therapies for HD with a reasonable duration and number of subjects. In conjunction with other recently described cognitive and affective ^10,42,43^ behavioral phenotypes in these animals, we propose that knock-in models provide greater face validity, and thereby potential translatability, than transgenic models of HD for future preclinical studies.

## Methods (1,500 words)

### Mice

We used 4 cohorts (detailed below) of mixed-gender B6.*Htt*^*Q111/+*^ mice for the described studies (Research Resource Identifier:IMSR\_JAX:003456). The creation of the *Htt*^*Q111/+*^ line has been described ^5^. Experiments for each cohort were approved by local institutional review in accordance with NIH or UK Animals Act animal care guidelines.

#### Cohort 1 (RNA sequencing)

The cohort used to generate the RNASeq data presented in Figure 1 was generated at Massachusetts General Hospital (MGH). Cohorts of 58 total mixed-gender littermate mice (12 x 3-month and 16 x 9-month *Htt*^*Q111/+*^; 12 x 3-month and 16 x 9-month *Htt^+/+^*) mice were generated by crossing B6.*Htt*^*Q111/+*^ males with C57Bl/6J females. The size of the *Htt* CAG tract ranged from 126 to 135 (mean 131), with no significant differences observed across the groups of this cohort. The mice were part of a larger study of the impact of dietary fat on peripheral metabolism - the mice described in Figure 1 were fed chow from weaning until 9 months of age with either 60% kcal/fat (Open Source Diet D12492) or 45% kcal/fat (Open Source Diet D1245). Striatal and cerebellar transcriptional effects of these diets were collapsed for the analysis presented in Figure 1.

#### Cohort 2 (Molecular natural history)

The cohort used to generate the molecular natural history data, presented in figures 2-5 and 7, was bred and maintained at the Jackson Labs (Bar Harbor, Maine) and shipped to the Western Washington University (WWU) vivarium approximately 2 weeks before the endpoints under investigation. Three groups of mice were sacrificed at 3 (90 ± 4 days), 9 (295 ± 2 days) or 12 (369 ± 5 days) months of age (Table 1).

#### Cohort 3 (Behavioral cohort, WWU)

The cohort used to collect the behavior data at WWU (porsolt swim test, elevated plus maze, light/dark box, sucrose solution preference) were bred and maintained at Jackson Labs (Bar Harbor, Maine) and consisted 40 female mice (20 *Htt^+/+^*; 20 *Htt*^*Q111/+*^). Htt CAG tract ranged from 107 to 119 (mean 114). Mice were shipped to the WWU vivarium at 8 months of age and acclimated to reversed light cycle conditions (lights on from 12am to 12 pm) for four weeks prior to testing at 9 months of age. Behavioral assays were conducted between 1:00 pm and 5:00 pm during the active phase.

#### Cohort 4: (Behavioral cohort, Cardiff)

The Cardiff cohort consisted of 16 mice (8 *Htt^+/+^*; 8 *Htt*^*Q111/+*^), bred in-house from founders originally obtained from Jackson Labs (Bar Harbor, Maine), was used for the FR1 and sucrose pellet preference experiments. The mice were housed under 12 hour light/dark cycle with free access to food and water outside of experimental periods. Experiments were run between the ages of 4 and 7 months of age. All experimental procedures were initiated between 8am and 9am.

### Tissue isolation and sample processing

After a three hour fast, mice in the molecular natural history cohort were euthanized via IP sodium pentobarbital injection (Fatal Plus, Henry Schein). Whole blood was collected by cardiac puncture with EDTA (25 μM) and plasma was extracted. Mice were then transcardially perfused with phosphate buffered saline (PBS) to clear tissues of blood. Whole brains were extracted and placed in a Brain Slicer Matrix (Zivic). A midline longitudinal cut was made to separate hemispheres. Coronal cuts were made in the left hemisphere 1- and 4-mm posterior to the junction between the olfactory bulb and cortex, resulting in a 3-mm thick corticostriatal block of tissue (Figure S2) that was formalin fixed (6-8 hours), paraffin embedded, sectioned (5-μm), and mounted onto glass slides for immunohistochemistry. The contralateral hemisphere was dissected into striatum, cortex, and cerebellum, and flash frozen for molecular analyses.

### Plasma chemistry and immune profiling

Plasma clinical chemistry was measured using an AU2700 Chemistry Analyzer (Beckman Coulter) at Phoenix Central Laboratories (Mukilteo, WA, USA). A panel of 37 cytokines, chemokines and acute phase reactants was assessed using a multiplexed cytometric bead array immunoassay (Mouse InflammationMAP 1.0, Myriad RBM, Inc., Austin, TX, USA).

### Library construction, RNA Sequencing and RNASeq analysis

For the RNA Sequencing studies in figure 1, RNA was extracted using the RNeasy Lipid Tissue Mini Kit (Qiagen). Quality control for RNA was conducted using the Agilent 2100 Bioanalyzer with Agilent RNA 6000 Nano kit. Libraries for sequencing were constructed using the Illumina TruSeq RNA Sample Prep Kit and sequenced on a HiSeq 2000 (2x50bp) to a read depth of 2.4 x 10^7^ ± 6.2 × 10^6^ per sample. The fastq files were aligned using the default parameters of SNAPR ^44^ (https://github.com/PriceLab/snapr) against the GRCh38 genome assembly, along with the transcriptome assembly gtf file from Ensembl, GRCh38.75. SNAPR generates read counts for both genes and transcripts simultaneously with alignment. All alignments were performed on Amazon EC2 c3.8xlarge instance using a Ubuntu14.04 base AMI. Differential gene expression was conducted using the edgeR package ^45^, and pathway enrichment using the HTSanalyzeR ^46^ in Bioconductor ^46^.

### mRNA quantification – QRT-PCR

Total RNA was extracted from the right hemi-striatum using the RNeasy Lipid Tissue Mini Kit (Qiagen). Tissue was homogenized in 500 µL of Qiazol Lysis Reagent for one minute at 4,000 rpm using a homogenizer (BeadBug; Benchmark Scientific). Aqueous and organic phases were separated by the addition of 100 µL of chloroform. All subsequent steps in were performed according to the manufacturer’s protocol. Reverse transcription was performed using the Superscript III First Strand Synthesis System (Life Technologies) following the manufacturer’s protocol. Quantitative real-time polymerase chain reaction (QRT-PCR) was conducted with the resulting cDNA using the Real-Time StepOne System (Applied Biosystems). Transcripts were analyzed by the relative quantitation standard curve method and normalized to β-Actin transcript levels. All taqman probes were purchased from Life Technologies and were as follows: Drd1a: Mm02620146\_s1, Drd2: Mm00438545\_m1, Cnr1: Mm01212171\_s1, Darpp32: Mm00454892\_m1, Penk: Mm01212875\_m1, N4bp2: Mm01208882\_m1, Islr2: Mm00623260\_s1, Pde10a: Mm00449329\_m1, H60b: Mm04243254\_m1, and Scn4b: Mm01175562\_m1.

### Immunohistochemistry

Deparaffinization washes were as follows: 2 x 3 min xylenes, 3 min xylenes:ethanol (1:1), 3 min 95% ethanol, 3 min 80% ethanol, 3 min 70% ethanol, 3 min 50% ethanol, H_2_O rinse. Heat-mediated antigen retrieval: 20 min at 95°C in citrate buffer (pH 6) or Tris-EDTA buffer (pH 9). Sections were blocked in 20% goat serum in PBS for 1 hour, followed by overnight incubation in primary antibody at 4°C; GFAP-stained slides were blocked with a mouse-on-mouse kit (Vector). Following wash, secondary antibody incubation was 1 hour at room temperature. Slides were mounted and sealed with DAPI fluormount-G (Southern Biotech). AIF1 slides were prepared from free-floating 40-µm coronal sections. Primary antibodies: mouse anti-aggregated HTT (1:750; DSHB; MW8), mouse anti-SQSTM1/p62 (1:300; Abcam; AB56416), rabbit anti-NeuN (1:750; Millipore; ABN78), mouse anti-DARPP32 (1:250; SCBT; H-3; sc-271111), rabbit anti-AIF1 (1:500; Wako), mouse anti-GFAP (1:500; Millipore; MAB3402). Secondary antibodies: Alexa 488 anti-mouse (1:1000; Life Technologies), Alexa 568 anti-rabbit (1:1000; Life Technologies).

### Image acquisition and analysis

For all immunolabeled sections, images were acquired with an IX-81 laser-scanning confocal microscope with Fluoview 1000 software (Olympus). Acquisition parameters were set such that a section with no primary antibody emitted no fluorescent signal. Multi-plane z-stack numbers and thickness were kept consistent for each set of sections. Maximum z-projections were compiled using ImageJ ^47^, and bit-depth was set to maximum range from 0-4095. NeuN and/or DAPI masks were used to measure neuronal and/or nuclear expression of MW8, p62, and DARPP32 (Figure 5B). AIF1 and GFAP were manually counted. Synaptophysin was quantified as percent area of image with synaptophysin immunoreactivity.

### Behavior

#### Elevated Plus Maze

The plus maze was made of white acrylic with two open arms (25x5x0.5; room lux ~475) and two closed arms (25x5x16cm) extending from a central platform elevated 50 cm above the floor. Each session began by placing a mouse in the intersection of the arms with its head directed toward the closed arm located opposite of the experimenter and ended after 10 minutes of free exploration. Activity was video recorded and analyzed using EthoVision XT 8 (Noldus).

#### Light/Dark Exploration

The light/dark exploration box used in this experiment was constructed of acrylic plastic divided into a clear, brightly illuminated side (27x27x30 cm; room lux ~475) and a black, fully enclosed side (18x37x30 cm) separated by a 5.5x5.5 cm opening that permitted the animals to move freely between each compartment. Each trial began by placing a mouse in the illuminated section of the apparatus facing the entrance to the dark section, and ended after 10 minutes of free exploration. Activity was video recorded and analyzed using EthoVision XT 8 (Noldus).

#### Porsolt Swim Test

Mice were placed into a clear 8-quart bucket (28 cm tall, 22 cm in diameter) filled ¾ full with room temperature water. Each mouse was placed into the bucket for a single 6-minute session, and swimming activity was recorded using a camera mounted on a tripod oriented toward the side of the apparatus. Inactivity during the last 4 minutes of each session was assessed by two separate experimenters blind to genotype (interrater reliability = 0.89), and a single composite score was calculated for subsequent analyses.

#### Sucrose Consumption Test

The mice were given a sucrose eating challenge to determine whether free access to sucrose would produce different levels of consumption between the genotypes. For these sessions the mice were given 95% sucrose pellets (AIN-76A, TestDiet, Richmond, IN) for 10 hours per day for 4 days, and normal lab chow for 14 hours. The amount of sucrose and chow consumed was measured daily by weight of 5 mg sucrose pellets/chow consumed and expressed as g/kg of bodyweight.

#### Operant Testing

Operant testing was conducted In 16 aluminium/steel 9-hole box (14cm x 13.5cm x 13.5cm) operant chambers (Campden Instruments, UK). On the rear wall of each chamber situated 15mm from the grid floor, a horizontal 9-hole light array was fixed that had 9 response holes (11mm diameter and 2mm apart) that contained lights at the rear and infra-red (IR) sensors at the front, such that nose poke responses to light stimuli could be detected with an IR beam break. For the present experiments only the central response hole (hole 5 of 9) was used with the other holes blocked. On the inner of the front wall a food magazine was placed that allowed the mouse to recover sweet liquid rewards (Yazoo strawberry milk, Campina Ltd, UK), delivered by peristaltic pump after a successful response to the light stimuli. Initial training consisted of non-contingent reward presentations to the magazine that were signalled by the illumination of a light in the magazine. Removal of the head from the magazine was detected by an IR beam across the magazine entrance which then reset the trail. This was followed by training the mice to respond with a nose poke to illuminated hole 5 on the light array. A response in lit hole 5 extinguished the light and illuminated the magazine light to signal reward delivery. On removal of the head from the magazine, the magazine light was extinguished and after a 2 s intertrial interval, a new trial began with the illumination of the stimulus light in hole 5 thereby continuing the fixed ratio 1 (FR1) schedule of reinforcement. Once all mice could successfully complete 30 trials in a 30 minute session, the reward seeking probes began. Nine 45 minute FR1 sessions were run using, 3 (x3) counterbalanced reward pump durations (40 ms, 60 ms, 100 ms), to determine sensitivity to different reward magnitudes with the key outcome measure being number of rewards obtained.

### Statistical Analysis

Statistics were processed in R 3.2.3 ^48^. Simulated distributions for power analysis were constructed with the mvrnorm function in the MASS package ^49,50^. The elastic net classifier was constructed using glmnet ^49^. Endpoint QRT-PCR and IHC data and code for power analysis are available at https://github.com/seth-ament/hd_endpoints. Graphics were produced using ggplot2 ^51^ and Illustrator (Adobe).

## Acknowledgements

CHDI Foundation grant A-8339 to J.B.C and NIH grant NS049206 to V.C.W.

## Author Contribution Statement

R.M.B. and S.R.C. completed a majority of the work (tissue collection, tissue processing, data analysis, manuscript preparation) and should be considered co-first authors. J.P.C. assisted with histological analyses. R.M.W. with M.E.M. prepared the RNAseq tissue. C.F., R.M.W., and N.D.P. designed analyses, aligned and analyzed the RNAseq data. S.P.B. and S.B.D. designed, ran and analysed the operant experiments and sucrose consumption task. D.S., E.W., B.S., L.J., A.G., J. A., M. A. conducted behavioral experiments including elevated plus maze, forced swim test, sucrose preference and light/dark box under the supervision and direction of J.P.C. and S.M. V.C.W assisted with the conception and design of studies and analytic approaches J.B.C. conceived the experiments and wrote the manuscript. All authors have seen and approved the manuscript in its final form.

## Additional information

The authors declare that the research was conducted in the absence of any commercial or financial relationships that could be construed as a potential conflict of interest.

## References

1. A novel gene containing a trinucleotide repeat that is expanded and unstable on Huntington’s disease chromosomes. The Huntington’s Disease Collaborative Research Group. Cell 72, 971–983 (1993).

2. Takano, H. & Gusella, J. F. The predominantly HEAT-like motif structure of huntingtin and its association and coincident nuclear entry with dorsal, an NF-kB/Rel/dorsal family transcription factor. BMC Neurosci. 3, 15 (2002).

3. White, J. K. et al. Huntingtin is required for neurogenesis and is not impaired by the Huntington’s disease CAG expansion. Nat. Genet. 17, 404–410 (1997).

4. Pouladi, M. A., Jennifer Morton, A. & Hayden, M. R. Choosing an animal model for the study of Huntington’s disease. Nat. Rev. Neurosci. 14, 708–721 (2013).

5. Wheeler, V. C. et al. Length-dependent gametic CAG repeat instability in the Huntington’s disease knock-in mouse. Hum. Mol. Genet. 8, 115–122 (1999).

6. Langfelder, P. et al. Integrated genomics and proteomics define huntingtin CAG length-dependent networks in mice. Nat. Neurosci. 19, 623–633 (2016).

7. Alexandrov, V. et al. Large-scale phenome analysis defines a behavioral signature for Huntington’s disease genotype in mice. Nat. Biotechnol. 34, 838–844 (2016).

8. Langbehn, D. R., Hayden, M. R., Paulsen, J. S. & PREDICT-HD Investigators of the Huntington Study Group. CAG-repeat length and the age of onset in Huntington disease (HD): a review and validation study of statistical approaches. Am. J. Med. Genet. B Neuropsychiatr. Genet. 153B, 397–408 (2010).

9. Menalled, L. et al. Systematic behavioral evaluation of Huntington’s disease transgenic and knock-in mouse models. Neurobiol. Dis. 35, 319–336 (2009).

10. Hölter, S. M. et al. A broad phenotypic screen identifies novel phenotypes driven by a single mutant allele in Huntington’s disease CAG knock-in mice. PLoS One 8, e80923 (2013).

11. Menalled, L. & Brunner, D. Animal models of Huntington’s disease for translation to the clinic: best practices. Mov. Disord. 29, 1375–1390 (2014).

12. Ross, C. A. & Tabrizi, S. J. Huntington’s disease: from molecular pathogenesis to clinical treatment. Lancet Neurol. 10, 83–98 (2011).

13. Sawiak, S. J., Wood, N. I., Williams, G. B., Morton, A. J. & Carpenter, T. A. Use of magnetic resonance imaging for anatomical phenotyping of the R6/2 mouse model of Huntington’s disease. Neurobiol. Dis. 33, 12–19 (2009).

14. Carroll, J. B. et al. Natural history of disease in the YAC128 mouse reveals a discrete signature of pathology in Huntington disease. Neurobiol. Dis. 43, 257–265 (2011).

15. Vonsattel, J.-P. et al. Neuropathological Classification of Huntington’s Disease. J. Neuropathol. Exp. Neurol. 44, 559–577 (1985).

16. Keum, J. W. et al. The HTT CAG-Expansion Mutation Determines Age at Death but Not Disease Duration in Huntington Disease. Am. J. Hum. Genet. 98, 287–298 (2016).

17. Wheeler, V. C. et al. Long glutamine tracts cause nuclear localization of a novel form of huntingtin in medium spiny striatal neurons in HdhQ92 and HdhQ111 knock-in mice. Hum. Mol. Genet. 9, 503–513 (2000).

18. Menalled, L. B. Knock-in mouse models of Huntington’s disease. NeuroRx 2, 465–470 (2005).

19. Björkqvist, M. et al. A novel pathogenic pathway of immune activation detectable before clinical onset in Huntington’s disease. J. Exp. Med. 205, 1869–1877 (2008).

20. Kwan, W. et al. Bone marrow transplantation confers modest benefits in mouse models of Huntington’s disease. J. Neurosci. 32, 133–142 (2012).

21. Landwehrmeyer, G. B. et al. Huntington’s disease gene: Regional and cellular expression in brain of normal and affected individuals. Ann. Neurol. 37, 218–230 (1995).

22. Li, S.-H. et al. Huntington’s disease gene (IT15) is widely expressed in human and rat tissues. Neuron 11, 985–993 (1993).

23. Strong, T. V. et al. Widespread expression of the human and rat Huntington’s disease gene in brain and nonneural tissues. Nat. Genet. 5, 259–265 (1993).

24. Rosas, H. D. et al. Evidence for more widespread cerebral pathology in early HD: An MRI-based morphometric analysis. Neurology 60, 1615–1620 (2003).

25. Falcon, S. & Gentleman, R. in Bioconductor Case Studies 207–220 (2008).

26. Subramanian, A. et al. Gene set enrichment analysis: a knowledge-based approach for interpreting genome-wide expression profiles. Proc. Natl. Acad. Sci. U. S. A. 102, 15545–15550 (2005).

27. Zwilling, D. et al. Kynurenine 3-monooxygenase inhibition in blood ameliorates neurodegeneration. Cell 145, 863–874 (2011).

28. Ko, J., Ou, S. & Patterson, P. H. New anti-huntingtin monoclonal antibodies: implications for huntingtin conformation and its binding proteins. Brain Res. Bull. 56, 319–329 (2001).

29. Mullen, R. J., Buck, C. R. & Smith, A. M. NeuN, a neuronal specific nuclear protein in vertebrates. Development 116, 201–211 (1992).

30. Wong, E. & Cuervo, A. M. Autophagy gone awry in neurodegenerative diseases. Nat. Neurosci. 13, 805–811 (2010).

31. Bjørkøy, G. et al. p62/SQSTM1 forms protein aggregates degraded by autophagy and has a protective effect on huntingtin-induced cell death. J. Cell Biol. 171, 603–614 (2005).

32. Walf, A. A. & Frye, C. A. The use of the elevated plus maze as an assay of anxiety-related behavior in rodents. Nat. Protoc. 2, 322–328 (2007).

33. Bourin, M., Michel, B. & Martine, H. The mouse light/dark box test. Eur. J. Pharmacol. 463, 55–65 (2003).

34. Mangiarini, L. et al. Exon 1 of the HD Gene with an Expanded CAG Repeat Is Sufficient to Cause a Progressive Neurological Phenotype in Transgenic Mice. Cell 87, 493–506 (1996).

35. Schilling, G. et al. Intranuclear inclusions and neuritic aggregates in transgenic mice expressing a mutant N-terminal fragment of huntingtin. Hum. Mol. Genet. 8, 397–407 (1999).

36. Van Raamsdonk, J. M. et al. Body weight is modulated by levels of full-length huntingtin. Hum. Mol. Genet. 15, 1513–1523 (2006).

37. Djousse, L. et al. Weight loss in early stage of Huntington’s disease. Neurology 59, 1325–1330 (2002).

38. Huntington Study Group & Huntington Study Group. Dosage effects of riluzole in Huntington’s disease: A multicenter placebo-controlled study. Neurology 61, 1551–1556 (2003).

39. Stout, J. C. et al. HD-CAB: A cognitive assessment battery for clinical trials in Huntington’s disease 1,2,3. Mov. Disord. 29, 1281–1288 (2014).

40. Thompson, J. C. et al. Longitudinal Evaluation of Neuropsychiatric Symptoms in Huntington’s Disease. J. Neuropsychiatry Clin. Neurosci. 24, 53–60 (2012).

41. Tabrizi, S. J. et al. Predictors of phenotypic progression and disease onset in premanifest and early-stage Huntington’s disease in the TRACK-HD study: analysis of 36-month observational data. Lancet Neurol. 12, 637–649 (2013).

42. Yhnell, E., Emma, Y., Dunnett, S. B. & Brooks, S. P. The utilisation of operant delayed matching and non-matching to position for probing cognitive flexibility and working memory in mouse models of Huntington’s disease. J. Neurosci. Methods 265, 72–80 (2016).

43. Oakeshott, S. et al. A mixed fixed ratio/progressive ratio procedure reveals an apathy phenotype in the BAC HD and the z_Q175 KI mouse models of Huntington’s disease. PLoS Curr. (2012). doi:10.1371/4f972cffe82c0

44. Magis, A. T., Funk, C. C. & Price, N. D. SNAPR: A Bioinformatics Pipeline for Efficient and Accurate RNA-Seq Alignment and Analysis. IEEE Life Sciences Letters 1, 22–25 (2015).

45. Robinson, M. D., McCarthy, D. J. & Smyth, G. K. edgeR: a Bioconductor package for differential expression analysis of digital gene expression data. Bioinformatics 26, 139–140 (2010).

46. Huber, W. et al. Orchestrating high-throughput genomic analysis with Bioconductor. Nat. Methods 12, 115–121 (2015).

47. Schindelin, J. et al. Fiji: an open-source platform for biological-image analysis. Nat. Methods 9, 676–682 (2012).

48. R Core Team. R: A language and environment for statistical computing. (2016).

49. Friedman, J., Hastie, T. & Tibshirani, R. Regularization Paths for Generalized Linear Models via Coordinate Descent. J. Stat. Softw. 33, 1–22 (2010).

50. Venables, W. N. & Ripley, B. D. Modern Applied Statistics with S. (2002).

51. Wickham, H. ggplot2: Elegant Graphics for Data Analysis. (Springer, 2016).

